# DSSP in Gromacs: tool for defining secondary structures of proteins in trajectories

**DOI:** 10.1101/2023.08.21.554196

**Authors:** S. V. Gorelov, A. I. Titov, O.A. Tolicheva, A. L. Konevega, A. V. Shvetsov

## Abstract

This work describes a fast implementation of software algorithm associated with determination of protein secondary structure based on the DSSP algorithm. This implementation is fully compatible with the DSSP v.4 algorithm and implemented as native Gromacs trajectory analysis module which allows to analyze molecular dynamics trajectories without any restrictions of the original DSSP implementation. Also this implementation works much faster then original DSSP v.4 algorithm.

**TOC Graphic:** 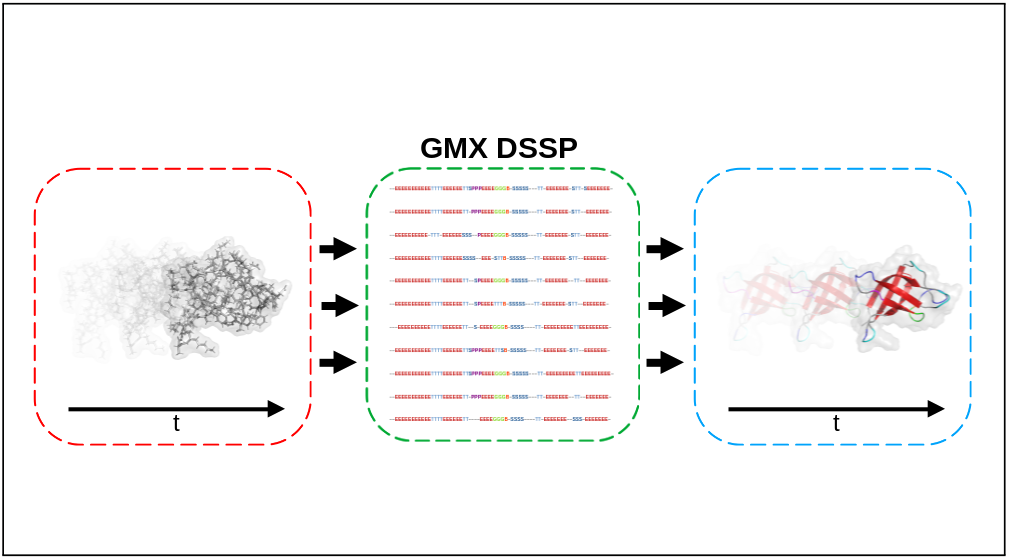

## Introduction

Hydrogen bonds provide most of the directional interactions that underlie protein folding, protein secondary structure, and molecular recognition, and thus are essential for understanding protein spatial organization and movement. The energy of such bonds is experimentally known, which varies greatly from *≈*5–6 kcal/mol for an isolated bond, to *≈*0.5–1.5 kcal/mol for proteins in solution. The cores of most protein structures consist of secondary structures such as *α*-helix and *β*-strand. This satisfies the hydrogen binding potential between the carbonyl oxygen of the main chain and the nitrogen of the amino group embedded in the hydrophobic core of the protein. Therefore, energetics and kinetics of the hydrogen bond must be optimal to allow fast selection and folding kinetics, conferring stability on the protein structure and providing the specificity required for selective macromolecular interactions. ^1,2^

To determine secondary structures, computational algorithms such as DSSP^3^ and STRIDE,^4^ are used. STRIDE is an algorithm for assigning elements of protein secondary structure taking into account protein atomic coordinates, determined using x-ray crystallography, nuclear-magnetic resonance or another method for determining the protein structure. In addition to hydrogen criteria used by a more common DSSP algorithm, the STRIDE appointment criteria also includes the dual-sided angle potentials. Thus, its criteria for determining individual secondary structure is more complex than that of DSSP. STRIDE energy function (see 1) has a member of the hydrogen bond containing the potential of Lennard-Jones 8-6 (see 2), which depends on the distance, and two angular planaries that reflect the optimized geometry of hydrogen connection (see 3 and 4):

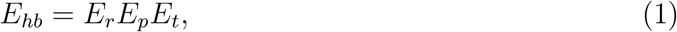

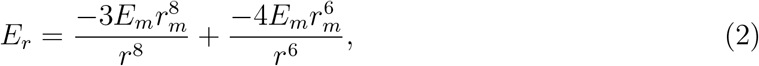

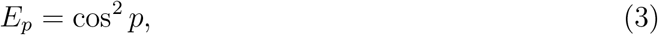

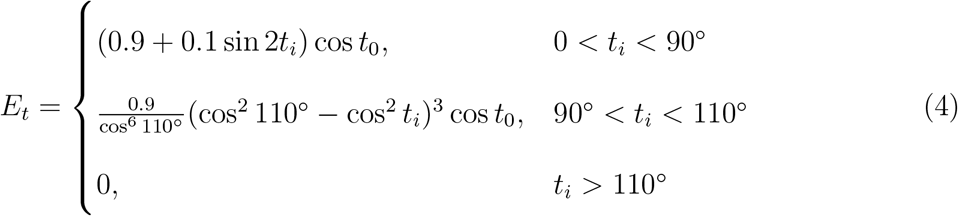

where *t*_*i*_ and *t*_0_ — angular deviations of the hydrogen atom from the bisector of an unexplored pair in the plane of an inconsistent pair of orbitals and from the plane of an inconsistent pair of orbitals, respectively, and *E*_*m*_ and *r*_*m*_ — optimal values for the hydrogen bond energy and length, respectively. For example, for hydrogen bonds N — H…O between atoms of the main protein circuit, values will be: *E*_*m*_ = −2.8 kcal/mol, *r*_*m*_ = 3.0 Å.

Criterias for individual secondary structural elements that are divided into the same groups as in DSSP also contain factors of statistical probability obtained as a result of empirical studies of resired structures with visually designated elements of the secondary structure extracted from the protein data bank.^4^

DSSP algorighm, on the other hand, uses coordinates of the structure to calculate hydrogen bonds arising in polypeptide between the elements of the main chain.^3^ Based on these connections, the DSSP algorithm searches for potentially possible secondary structures for a specific amino acid residue according to clear criteria for compliance with the template of secondary structures and assigns them to the amino acid. The final result is set according to the system of priorities of secondary structures. Thus, as a result of the algorithm, a full-fledged and unambiguous recording of secondary structure designations is formed for a separate protein.

The implementation of the DSSP algorithm in the Gromacs software package can be used to unambiguously determine the secondary structure of proteins using established hydrogen bond patterns. It is presented as a trajectory analysis module and, thus, able to process not only simple structures from the Protein Data Bank, but also large trajectories of simulated systems. The implementation combines the accuracy of DSSP v.4,^5^ wide variability due to the presence of additional functions and high speed of analysis.

## Implementation

To calculate the secondary structure using our implementation of the DSSP algorithm in Gromacs, firstly, it is necessary to obtain atomic coordinates of the atoms of the protein structure. The input is: a structural file, which explicitly specifies information about the coordinates and a topology file, which contains information about the composition of protein residues. The topology is analyzed first. It selects only the atoms of the main chain (C*α*, C, O, N and H atoms; denoted in Gromacs as “MainChain+H”) and stores their coordinates. Each residue is assigned a “previous” and a “next” residue, if any.

Next, molecular dynamics trajectory frame is analyzed. Each frame is processed independently. In it, the hydrogen bond energy is calculated between the selected protein residues. There are 2 methods for selecting desired residues for calculation. First one checks hydrogen bond energy between all residues in the protein according to the principle “each with each”, which is a complete analogy of the original DSSP v.4 algorithm.^5^ An alternative way is to use the built-in Gromacs function - Neighbor Search, which, using a box with periodic boundary conditions, quickly finds the neighbors of any residue in a certain range (the minimum, as well as recommended, range value for determining neighbors is 9 Å). Using Neighbor Search allows you to significantly speed up the process of finding nearby residues. Regardless of the chosen method, the presence of hydrogen bonds between the residues is checked each time. ^6^

### Hydrogen bond patterns

Hydrogen bonds in proteins have a small overlap of wave functions and are well described by the following electrostatic method.^3^ When 2 residues are compared with each other, then the energy criterion is calculated by the formula:

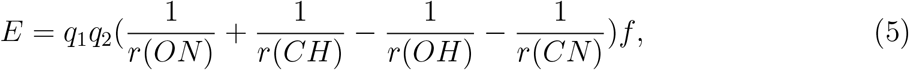

where *q*_1_ = 0.42*e, q*_2_ = 0.20*e, f* = 332, and *r*(*AB*) is the interatomic distance from atom *A*

to atom *B*.

It is generally accepted^3^ that a hydrogen bond exists between two residues if the energy of this bond does not exceed -0.5 kcal/mol. A good hydrogen bond has a binding energy of about -3 kcal/mol. The bond donor is the amino group, the acceptor is the carbonyl group. Each amino acid can be twice an acceptor and twice a hydrogen bond donor (except for proline, it cannot be a hydrogen bond donor because it lacks a hydrogen atom associated with nitrogen). As a result, for each residue there is a record of the 2 most energetically favorable hydrogen bonds (if any) and the corresponding energy values.^3^

Based on spatial arrangement of atoms and patterns of hydrogen bonds between residues, the secondary structure of a protein can be determined. If an amino acid does not have any features, it remains a loop and is designated as “*∼*”. If the distance between the C atom of the main chain of one residue and the N atom of the main chain of the residue following it exceeds 2.5 Å, then it is considered that there is a gap between them (which is not a secondary structure in the classical sense), and it is designated as “=“. Bends (“S”) are areas in the protein with a large curvature. The residue is considered to be a bend if the curvature of the chain in the central residue of five successive ones makes an angle of more than 70^*°*^, which is defined as the angle between the direction of the main chain of the first and last three residues of those considered.

There are three types of turns — 3-turn, 4-turn, 5-turn. All turns, regardless of their type, are designated as “T”. Each turn corresponds to helix of specific type: a 3-turn corresponds to a 3_10_-helix; A 4-turn corresponds to an *α*-helix; A 5-turn corresponds to a *π*-helix. These helices are respectively designated as “G”, “H” and “I”. Amino acid *i* is an n-turn if it has a hydrogen bond with *i* + n amino acid. Two successive turns of the same type form a helix of the corresponding type, along the edges of which there will be turns.

Two overlapping minimal helices (formed by the overlap of two turns of the corresponding type), displaced by two or three residues, are combined into one long helix. Such a combination allows one to take into account the fact that long helices may deviate from regularity in formation due to the absence of hydrogen bonds critical for the formation of a pattern of turns. Such imperfections are often associated with helix kinking, for example due to a proline residue. There are also unique polyproline turns (they don’t have a unique designation) and helices (polyproline II helices, also known as *κ*-helices; designated as “P”). They are defined somewhat differently: if the value of the torsion angles *φ* and *ψ* for amino acid *i* falls within a fixed range of values (6a and 6b, respectively), then it forms a polyproline turn.

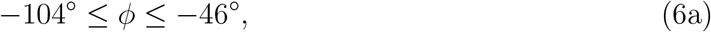

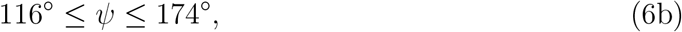

Depending on the value specified by the user in the “-ppstretch” parameter, 2 or 3 (default) consecutive polyproline turns form a polyproline II helix with polyproline turns at the ends.

There are also *β*-bridges (“B”) and *β*-strands (“E”). Two non-intersecting segments with three residues each, *i* - 1, *i, i* + 1 and *j* - 1, *j, j* + 1, form either a parallel or antiparallel *β*-bridge, depending on which of the two main patterns coincides (see Fig. 3). A *β*-strand consists of two *β*-bridges of the same type, connected by no more than one additional residue on one strand and no more than four additional residues on the other strand (see Fig. 4).

**Figure 1:**
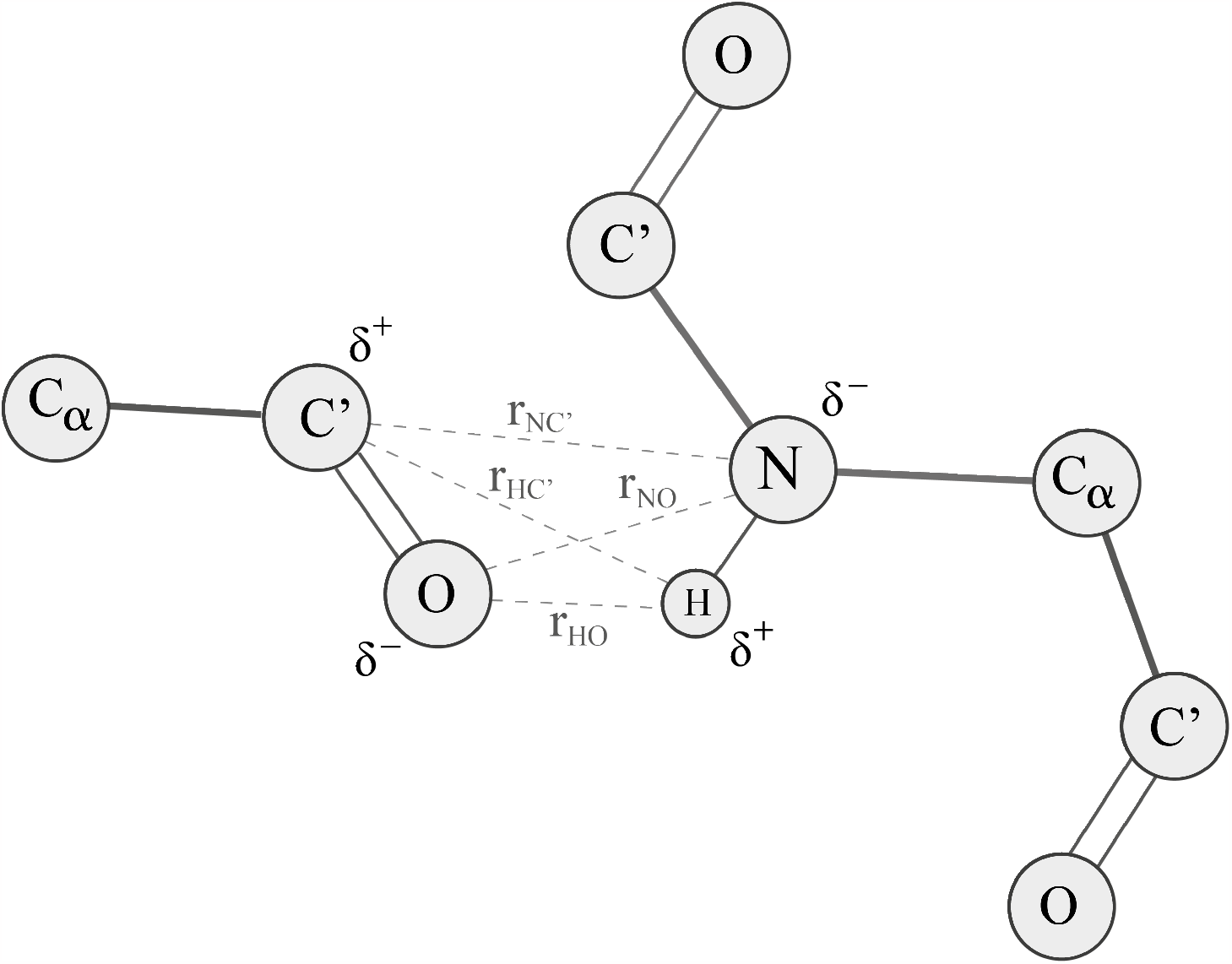
Schematic representation of a hydrogen bond formation between two residues.

**Figure 2:**
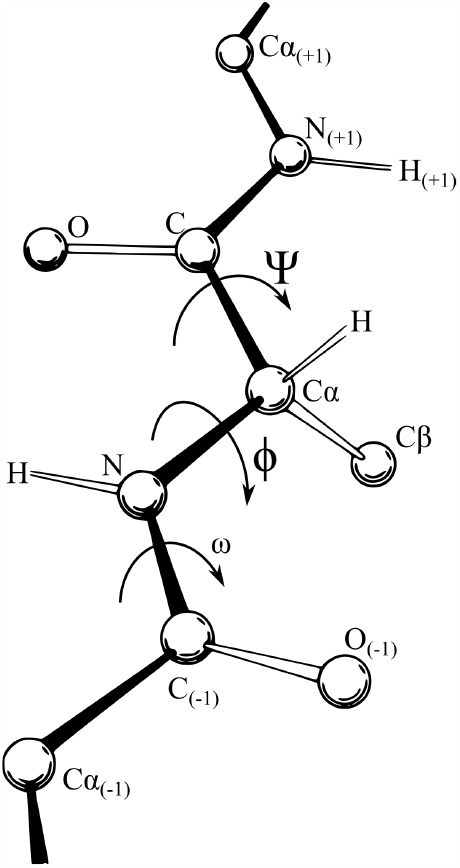
Schematic representation of the torsion angles *ϕ* and *ψ* relative to an arbitrary polypeptide residue.

**Figure 3:**
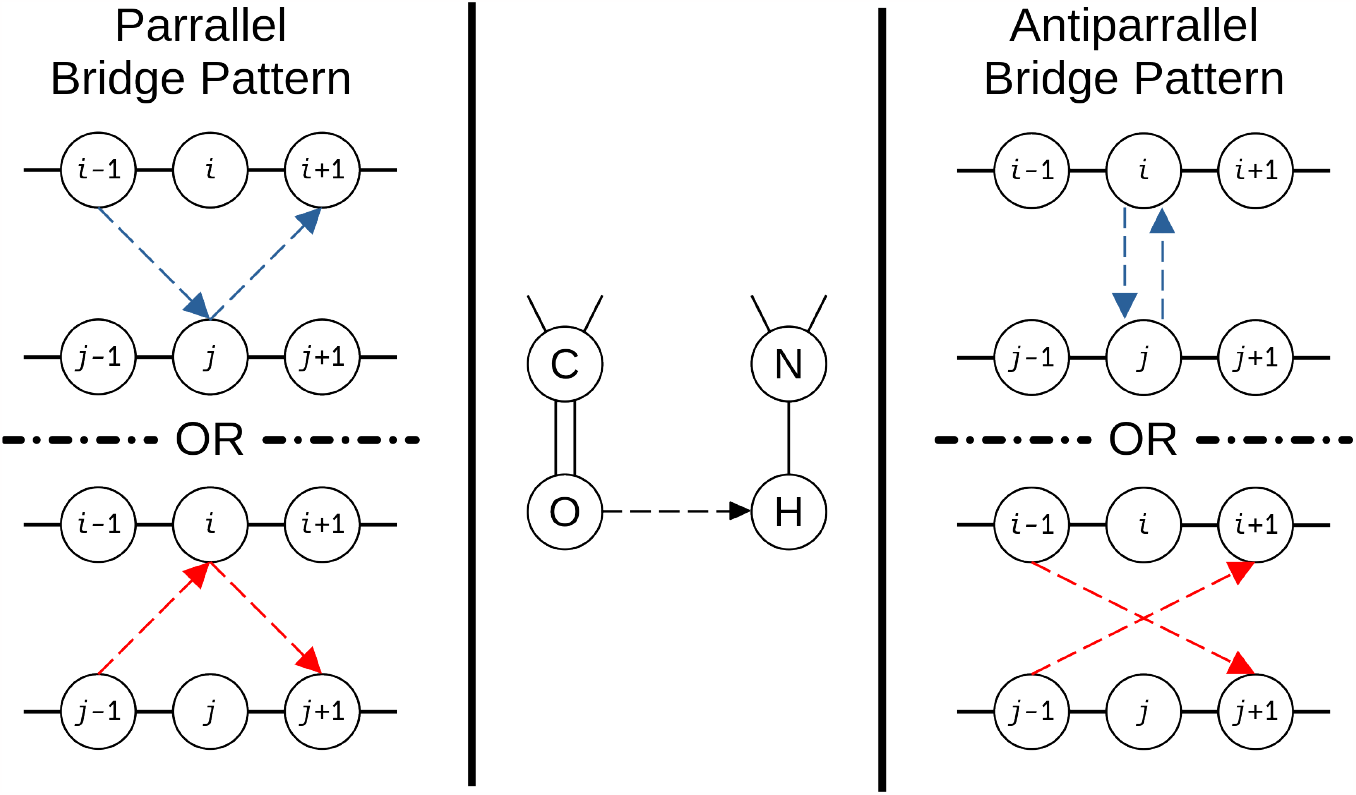
Schematic notation of *β*-bridge detection pattern.

**Figure 4:**
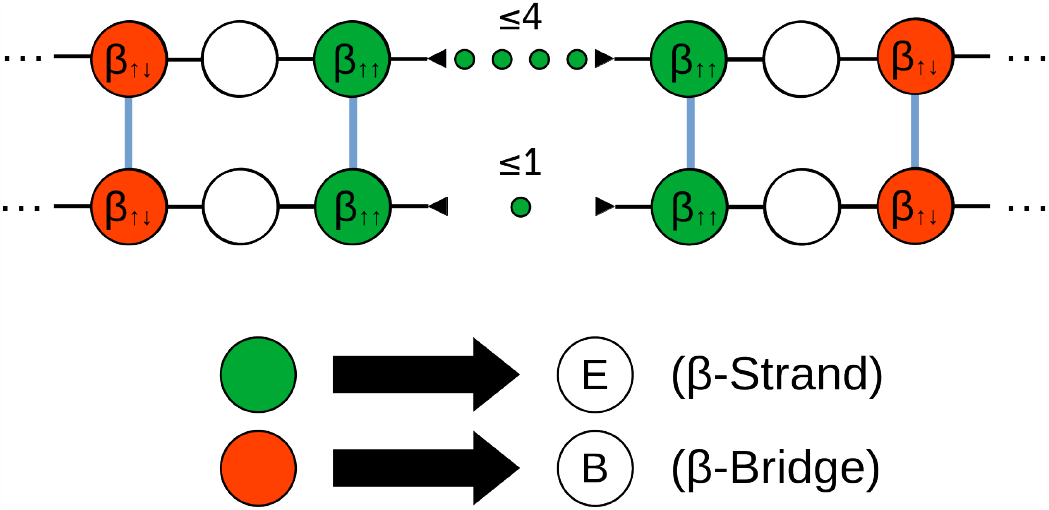
Schematic notation of *β*-bridge and *β*-strand assignment patterns.

**Figure 5:**
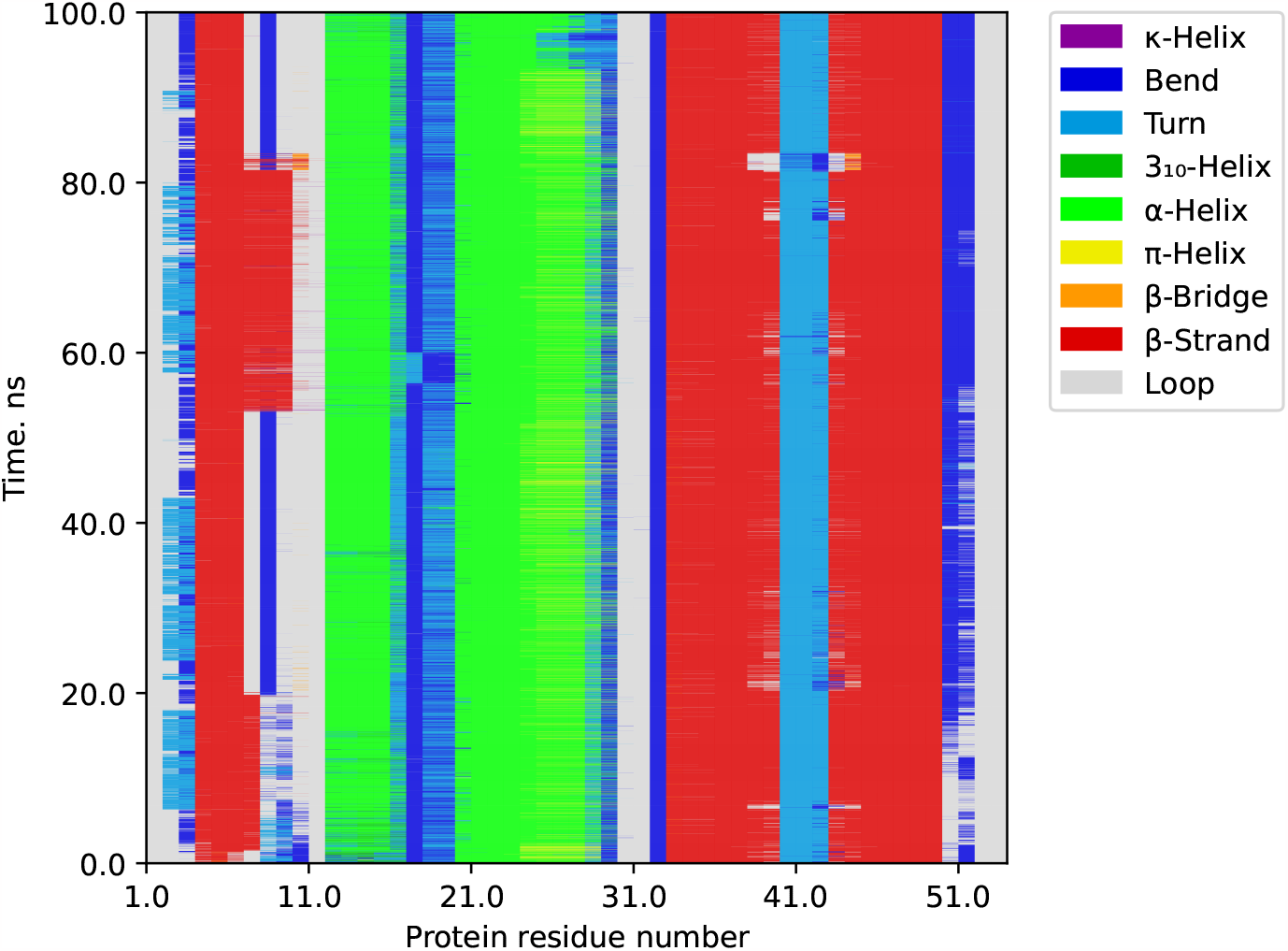
Map of secondary structures for the relative positions of residues in the defensin-like protein of *Pentadiplandra brazzeana* trajectory.

**Figure 6:**
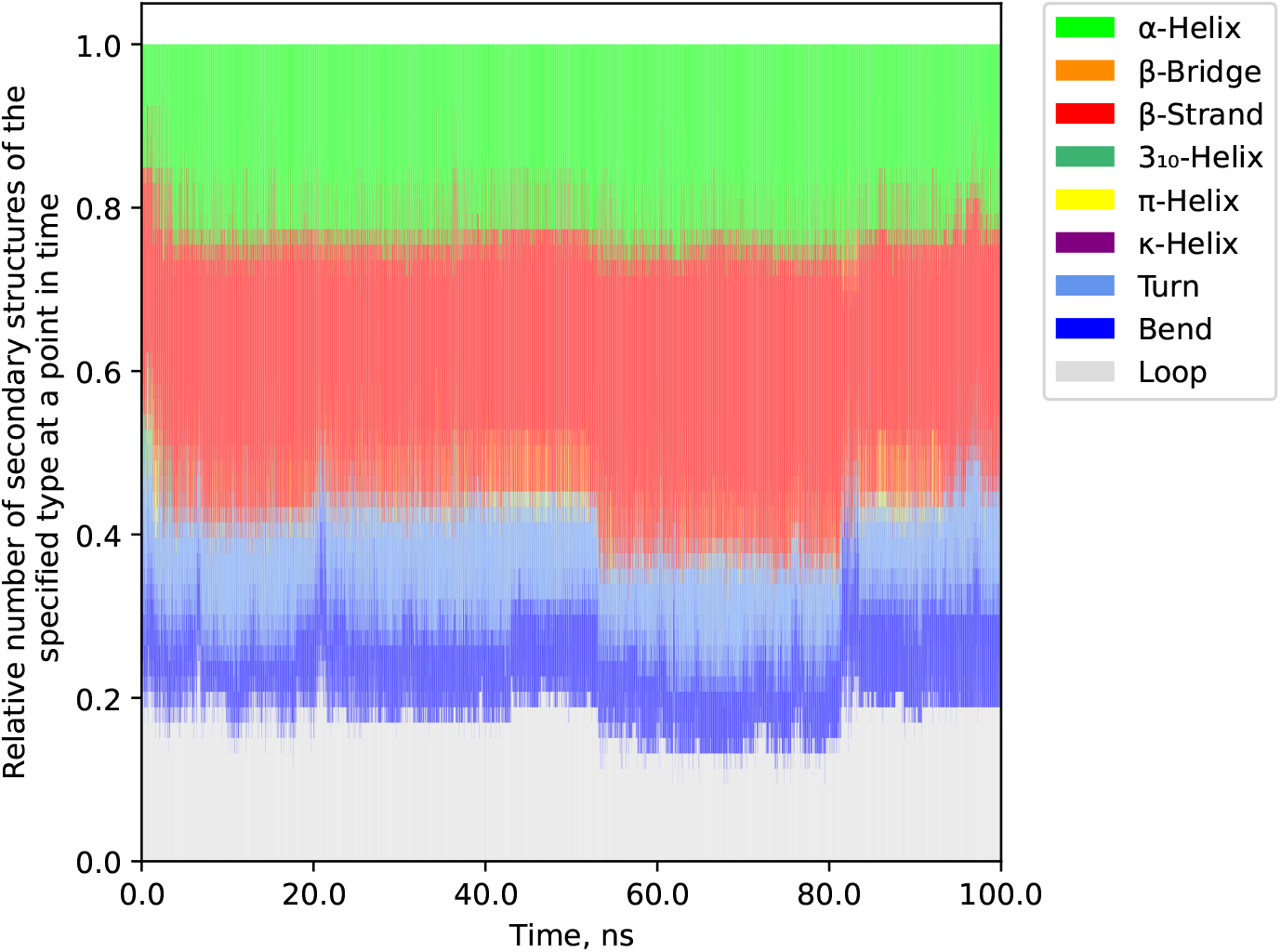
Graph of the relative number of secondary structures versus time in the trajectory of the defensin-like protein of *Pentadiplandra brazzeana*.

One amino acid can have multiple secondary structure assignments. But, since the output must always give defined values for each residue, a problem with assignment of secondary structure arises. To solve this problem, there is an unambiguous priority system similar to DSSP ver. 4: The structure with the higher priority will be selected. For example, the “P” polyproline II helix is not assigned to a residue if it already has some secondary structure, *β*-bridges are assigned only when the criterion for their existence is met and the residue is not a *β*-strand. As in the original DSSP algorithm (after version 2.1.0^7^), *π*-helices can take a precedence over *α*-helices assignments (corresponds to the “-pihelix” parameter), however, by entering the “–nopihelix” parameter, the user can shift the priority when setting the secondary structure to prioritize *α*-helices, which will correspond to the output of DSSP (prior version 2.1.0).

For convenience, you can look at table 1, which contains all of the above one-symbol designations of the secondary structures.

**Table 1:**
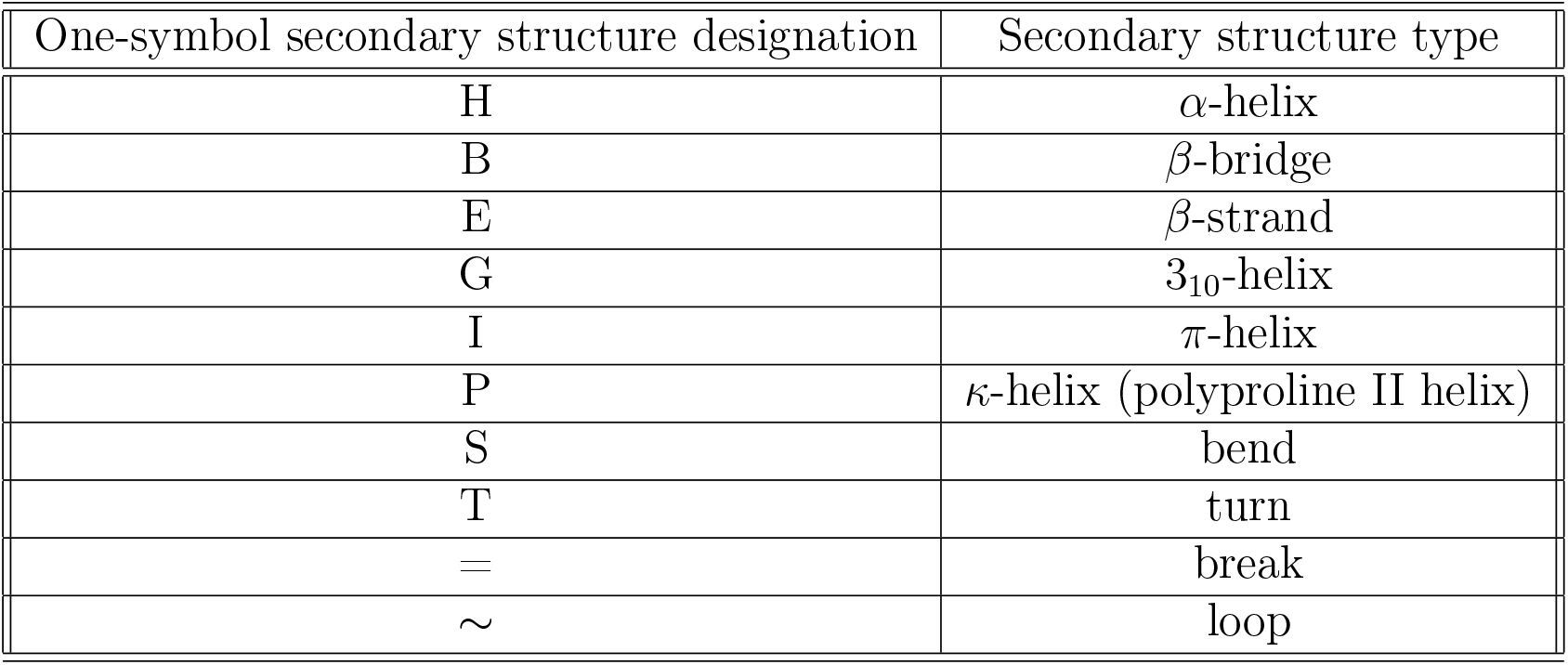
One-symbol secondary structure designations used in our implementation of DSSP algorithm.

| One-symbol secondary structure designation | Secondary structure type |
| --- | --- |
| H | α-helix |
| B | β-bridge |
| E | β-strand |
| G | $3_{10}$-helix |
| I | π-helix |
| P | κ-helix (polyproline II helix) |
| S | bend |
| T | turn |
| = | break |
| ∼ | loop |

### Additional parameters

It is important to take into account that a protein does not always have a hydrogen atom in its structure: for example, hydrogen atoms are difficult to detect by X-ray diffraction, so they usually do not exist in structures obtained by this method. In such case, along with the mode of using native hydrogens, the mode of creating hydrogen pseudo-atoms for each residue, except for proline (the “-hmode dssp” parameter) was implemented: for each residue, the hydrogen atom H is given the atomic coordinates of the nitrogen atom N of the same residue, after which the vector of the pseudo-atom H is added to the vector drawn from the O atom to the C atom of the previous residue:

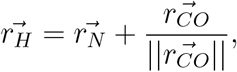

However, by default, it is assumed that all hydrogen atoms are present in the structure (parameter “-hmode gromacs”), since in formats native to Gromacs, hydrogen atoms are mandatory.

If one or more atoms of the main chain (except for hydrogen atoms, their presence/absence is controlled by the “-hmode” parameter) is missing from the structure, the algorithm will ignore these atoms when calculating angles and distances. However, specifically to improve the accuracy of calculations and the convergence of results with the original DSSP v.4 algorithm, the “-clear” parameter was implemented, which allows discarding entire protein residues during topology analysis if they are found to lack critically important atoms for analysis.

The developed DSSP algorithm also makes it possible to search for existing hydrogen bonds not only using the energy criterion (see 5), but also using the geometric criterion:

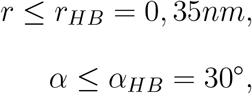

where *r* is the distance between donor and acceptor of hydrogen bond, and *α* is the angle between acceptor and hydrogen. The value *r*_*HB*_ = 0.35 nm corresponds to the first minimum of the radial distribution functions of water in the SPC/E model.^6^

To do this, the user must select one of the methods between “-hbond energy” and “-hbond geometry”. It should be noted that calculations using the geometric criterion determine a smaller number of hydrogen bonds between residues than calculations using the energy one. Consequently, a search by hydrogen bond patterns when calculating by a geometric criterion will find fewer grounds for an unambiguous determination of the secondary structure.

An option was developed to visualize the obtained results in the form of a graph of the number of secondary structures depending on time (per frame). The calculation is performed by entering the “-num” parameter by the user. The algorithm will select the number of received secondary structures of each unique type on each frame and form an output file from this data, from which it will be possible to plot.

### Example applications

As the result of this work, our own implementation of the DSSP algorithm was developed, which generates a sequence of characters in the output file (or several lines when a file with several frames is analyzed) denoting the secondary structure of the protein. It is important to note that this output file does not contain information about the specific time of existence of secondary structures in the protein, since the time between frames can be absolutely arbitrary and depends on the parameters specified by the user when generating the trajectory.

The developed algorithm was applied to the molecular dynamics trajectory of the defensin-like protein of *Pentadiplandra brazzeana* (Brazzein) pdbid:1BRZ.^8^ The duration of the trajectory is 1*μ*s.

## Discussion

Obtained results from the developed algorithm were compared with the results of the DSSP v.4 algorithm.^5^ Despite the greater variability of the developed algorithm, with certain given parameters, it allows the output of data that does not differ in content from the output of DSSP v.4. Thus, we can assume that the developed algorithm is not inferior in accuracy to the DSSP v.4 algorithm.

In the graphs above, it is shown that in Brazzein protein, residues 5-7 form a relatively stable *β*-strand, which over time can temporarily expand to a 5-10 *β*-strand. Residue 9 is stably a bend, but only if there is not expanded 5-7 *β*-strand in the structure. Residues 13-17 and 21-29 form relatively stable *α*-helices, between which bends with turns are formed. Moreover, the 13-17 helix is (initially) a 3_10_-helix that transforms into an *α*-helix over time. The end of the 21-29 *α*-helix (residues 25-29) sometimes turns into rare *π*-helices. Residues 34-39 and 44-50 form stable *β*-strands, between which turns usually form. Residues 2, 12, 31-32, 53-54 almost throughout the entire trajectory of molecular dynamics do not receive assignments of secondary structures. The secondary structures of the defensin-like protein of *Pentadiplandra brazzeana* determined by the developed algorithm coincide with the assignments of the secondary structures obtained experimentally.^9,10^

### Comparison of algorithms’ “run time”

The speed of the old and new implementations of the DSSP algorithm was tested on trajectories of different sizes for defensin-like protein of *Pentadiplandra brazzeana*, Sars-Cov-2 Non-Structural Protein 12, T7 RNA polymerase and scFv antibodies: 3B12, 3H10, B11-1, B11-2 and ST4. In all cases, to obtain records of secondary structures, only protein atoms were used as input.

It is important to note that the original DSSP algorithm is not capable of performing secondary structure analyzes for molecular dynamics trajectories. Therefore, to compare running time of said algorithms, pre 2023.1 DSSP implementation in Gromacs was used, which converts trajectory into a readable DSSP v.4 format (.pdb) and then launch analysis through DSSP v.4.

According to the data presented on figures 7, 8 and 9 in supporting information section, we can conclude that the developed implementation based on the DSSP algorithm works much faster than DSSP v.4 counterpart. This result was achieved due to the correct integration of the DSSP algorithm into the Gromacs trajectory analysis module.

**Figure 7:**
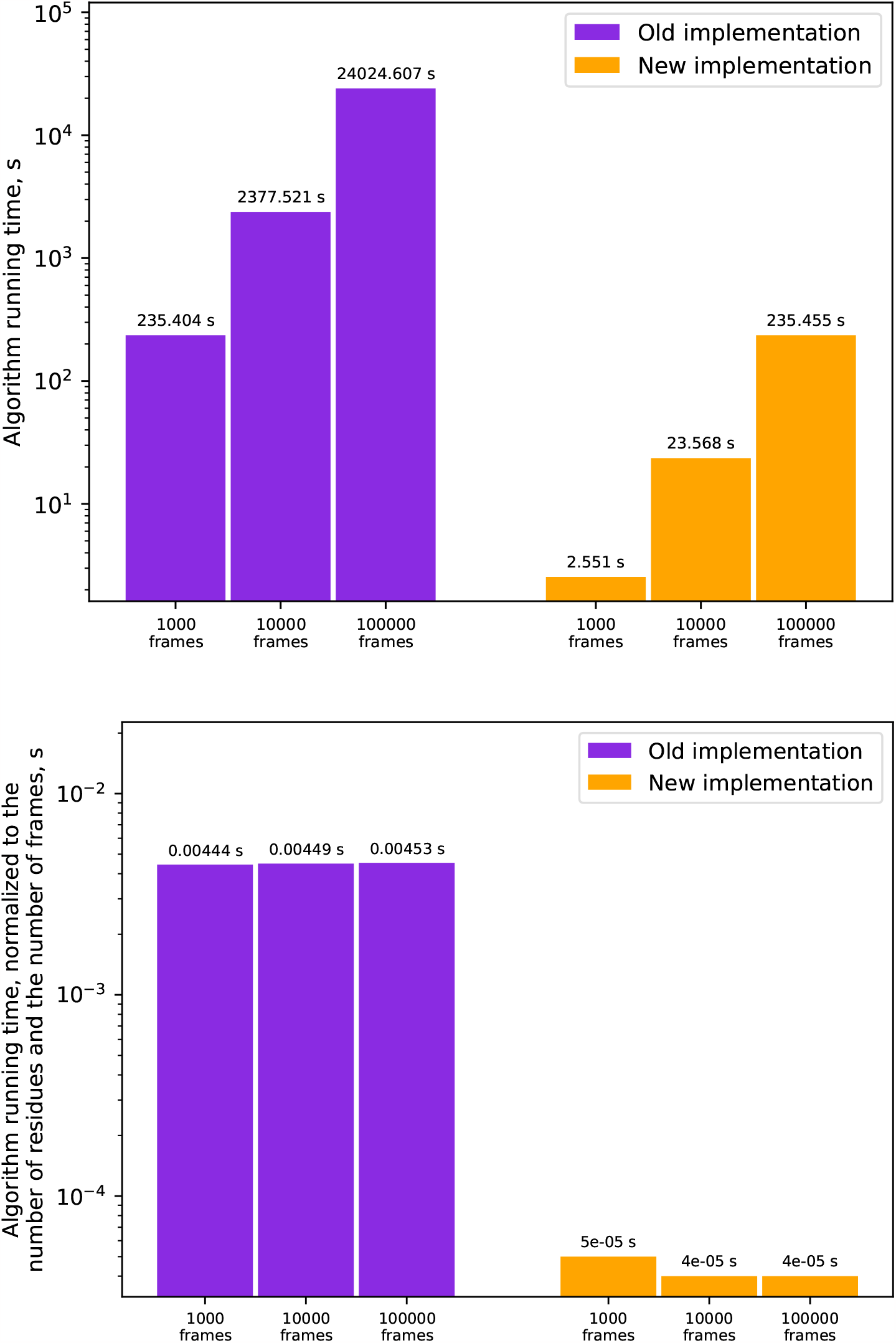
Comparison of the running time of the algorithms on the trajectories of the defensin-like protein of *Pentadiplandra brazzeana* (protein length 53 a.a)

**Figure 8:**
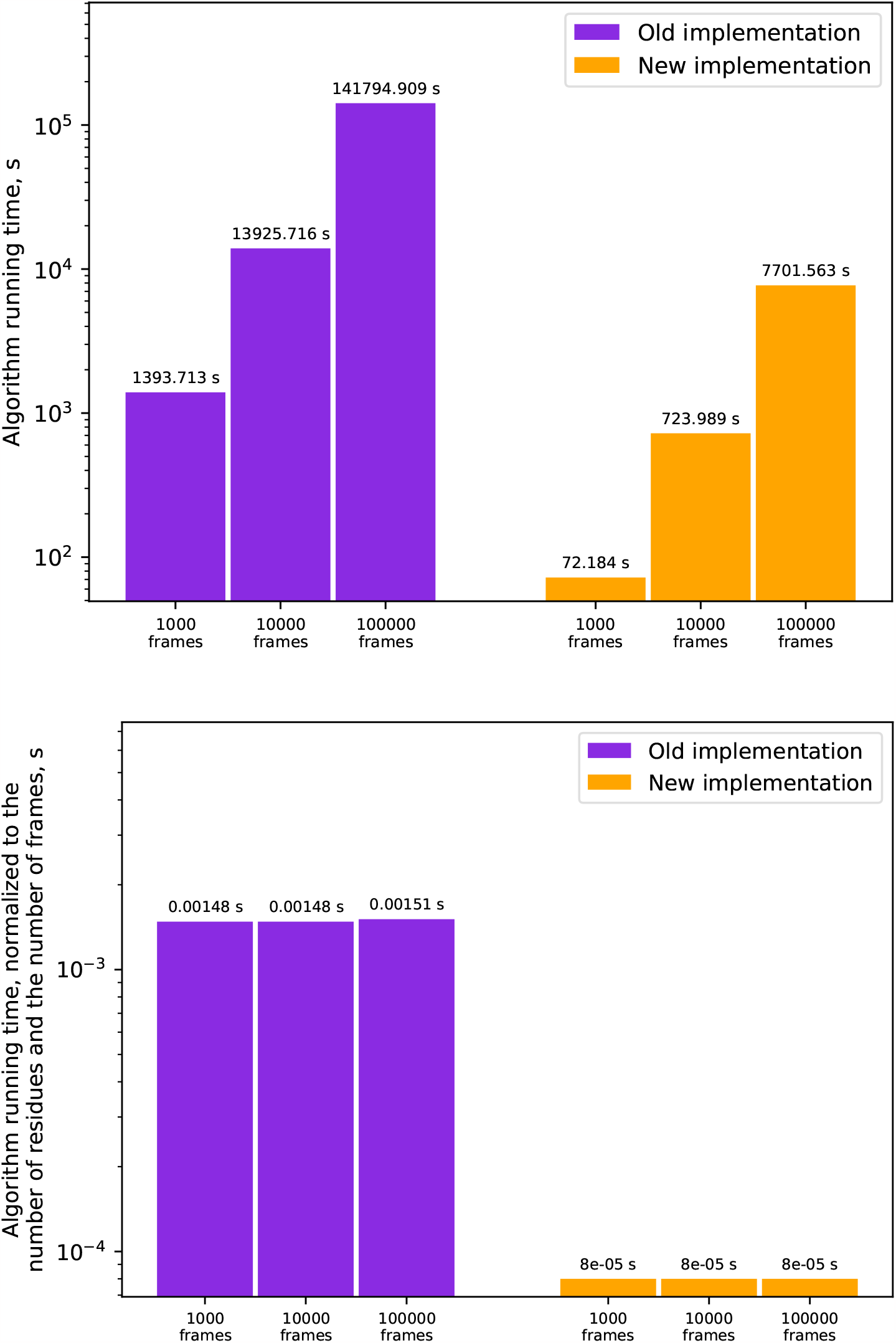
Comparison of the running time of the algorithms on the trajectories of the Sars-Cov-2 Non-Structural Protein 12 (protein length 941 a.a.)

**Figure 9:**
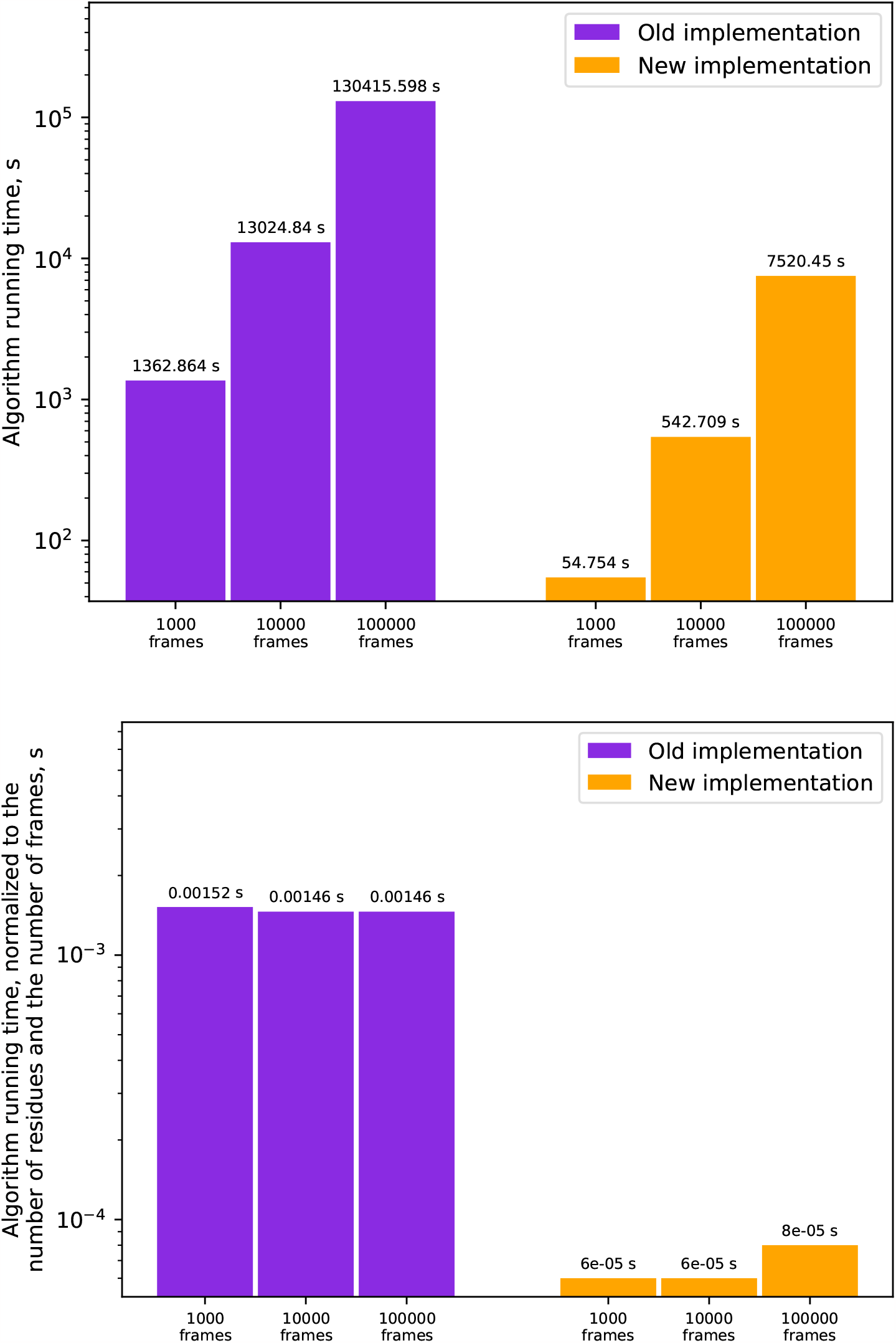
Comparison of the running time of the algorithms on the trajectories of the T7 RNA polymerase (protein length 894 a.a.)

**Figure 10:**
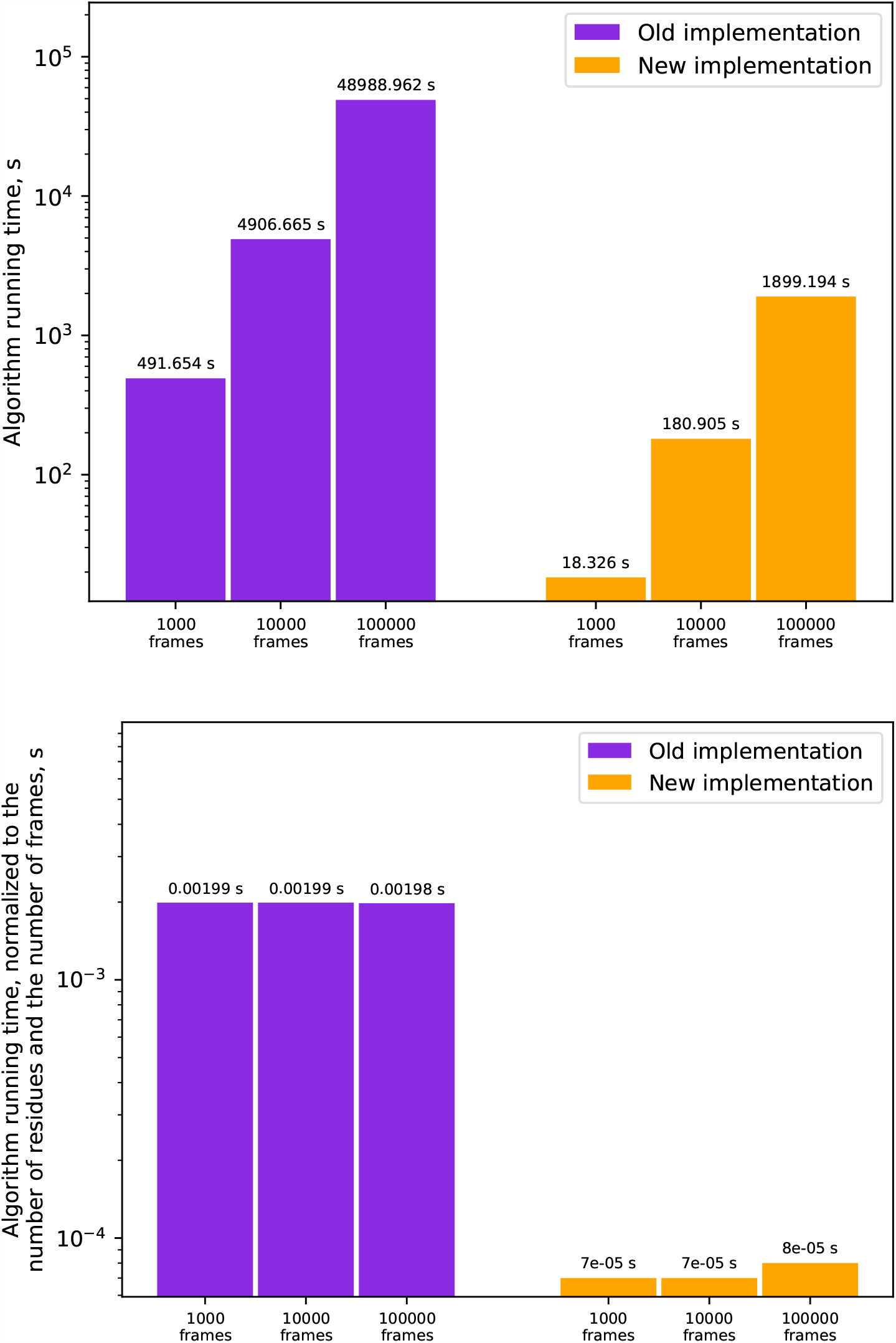
Comparison of the running time of the algorithms on the trajectories of the scFv antibody 3B12 (protein length 247 a.a.)

**Figure 11:**
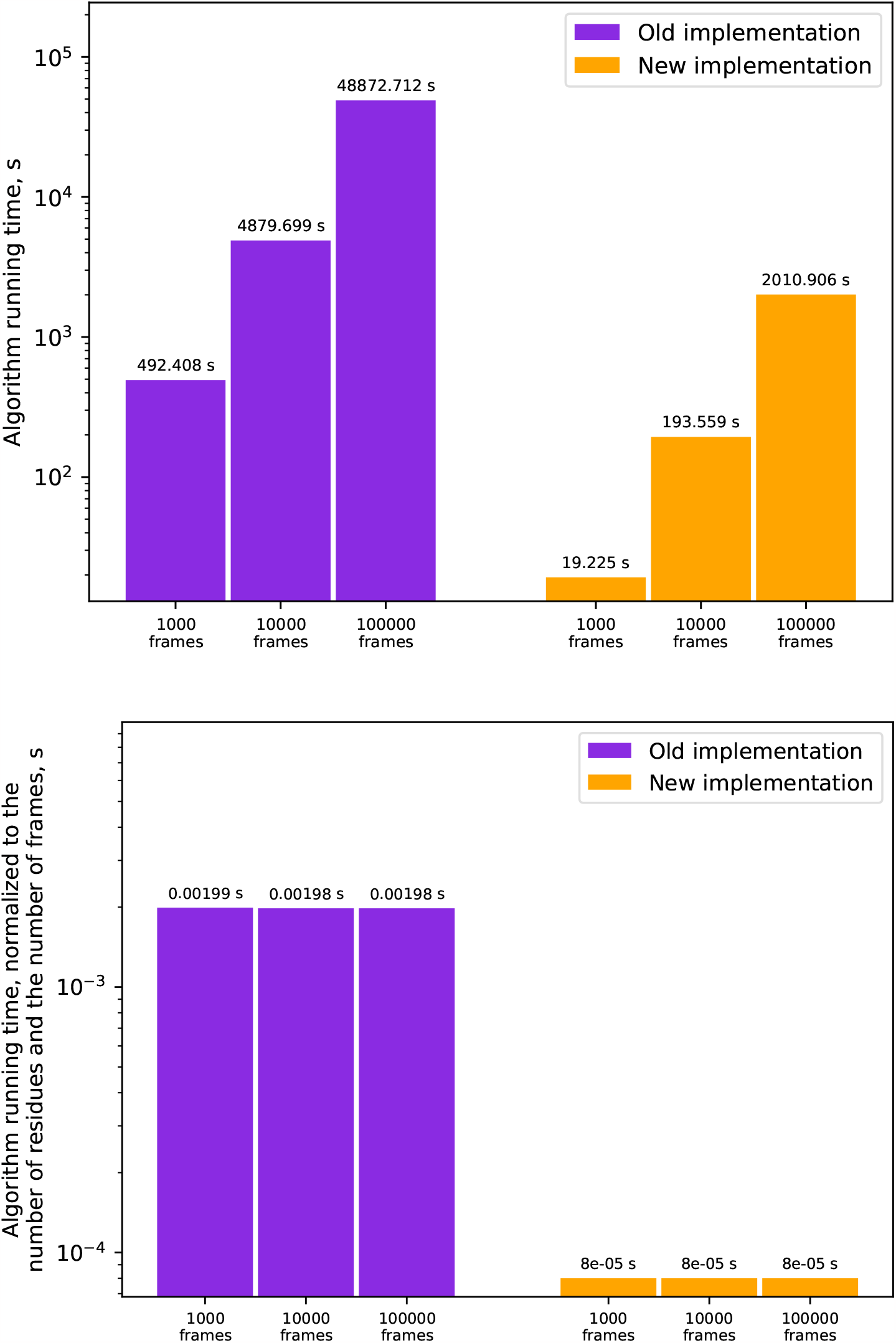
Comparison of the running time of the algorithms on the trajectories of the scFv antibody 3H10 (protein length 247 a.a.)

**Figure 12:**
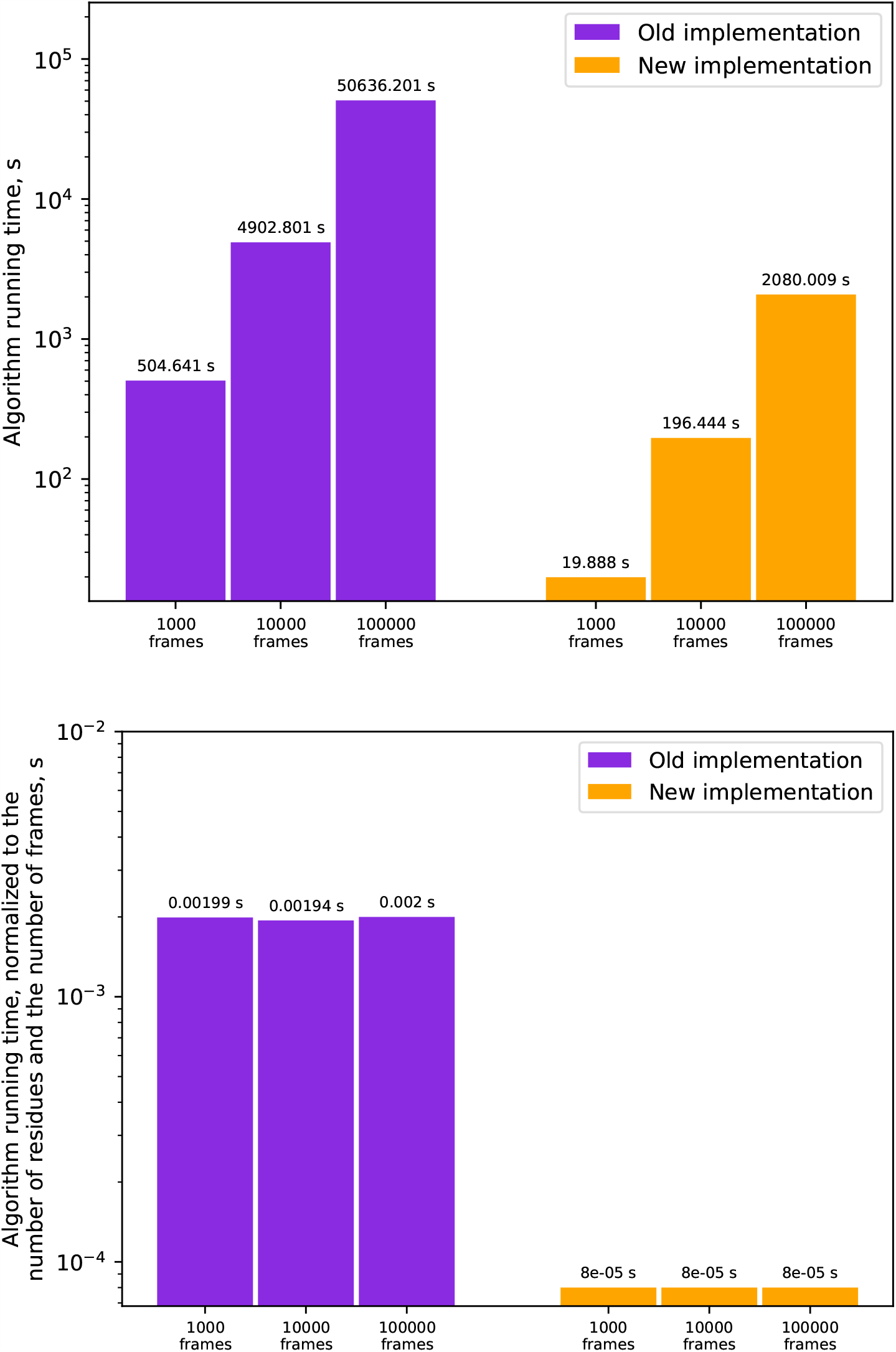
Comparison of the running time of the algorithms on the trajectories of the scFv antibody B11-1 (protein length 253 a.a.)

**Figure 13:**
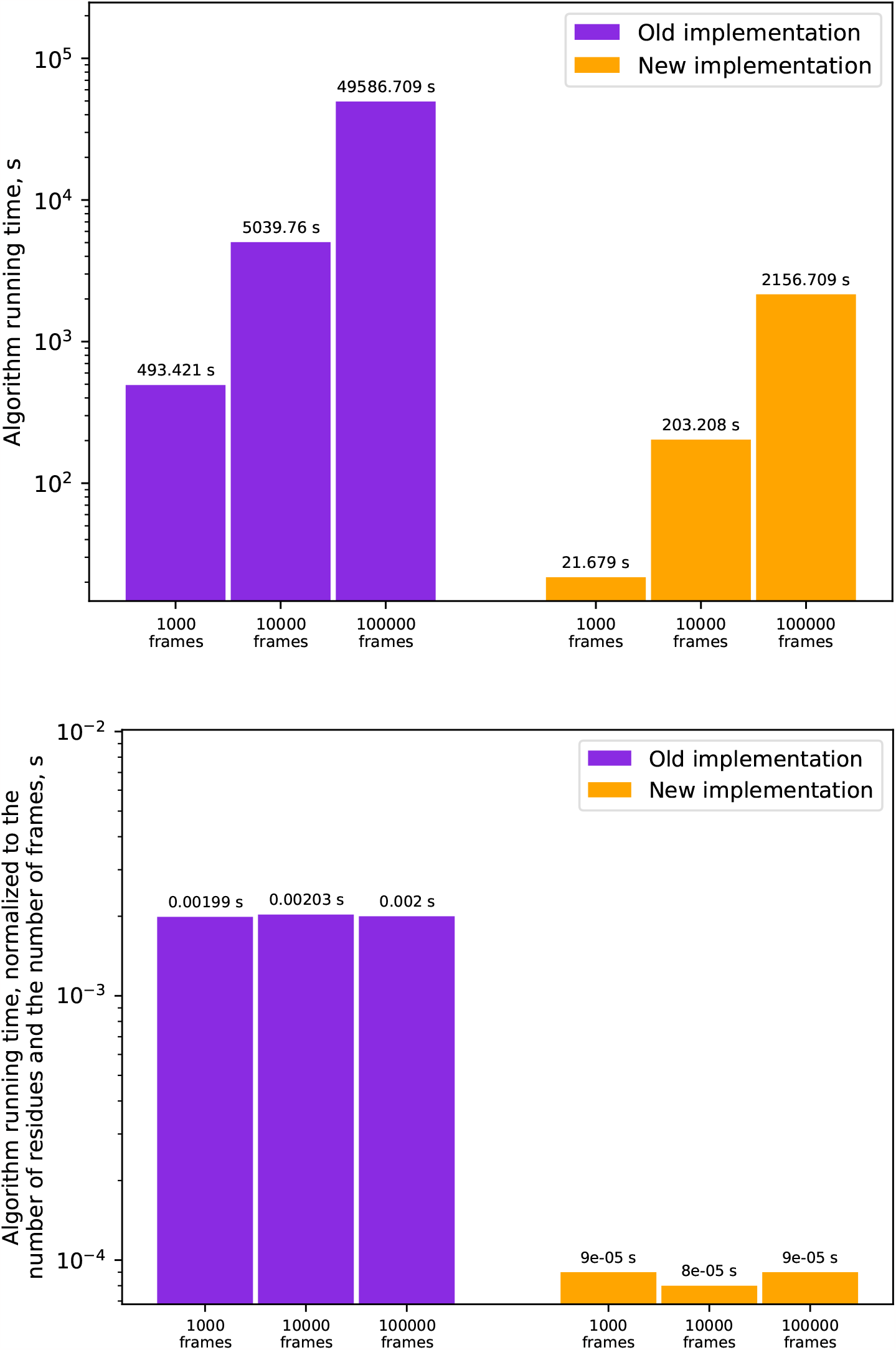
Comparison of the running time of the algorithms on the trajectories of the scFv antibody B11-2 (protein length 248 a.a.)

**Figure 14:**
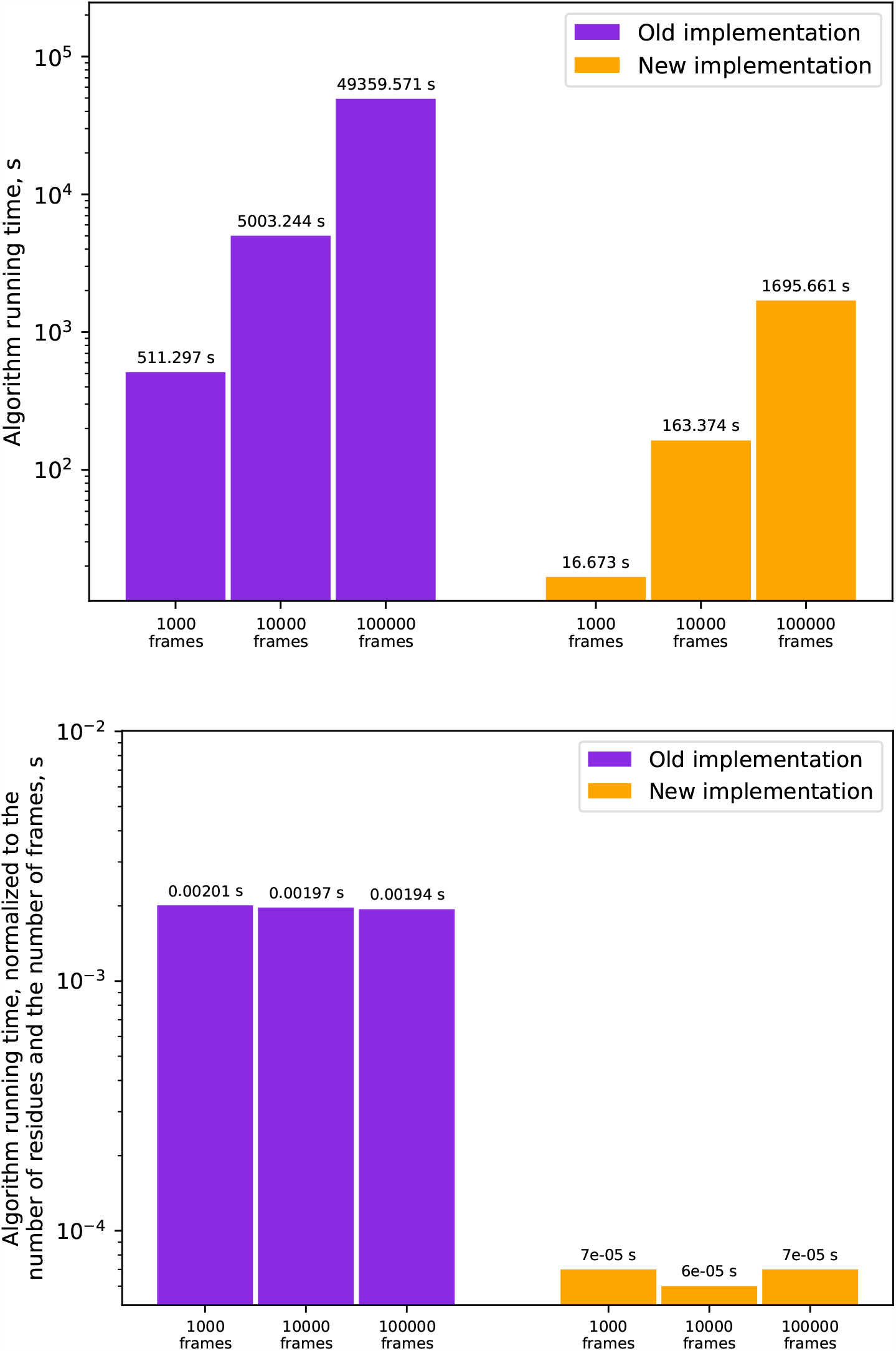
Comparison of the running time of the algorithms on the trajectories of the scFv antibody ST4 (protein length 254 a.a.)

## Conclusions

It is demonstrated that the developed algorithm for determining secondary structures is better than the original DSSP v.4^5^ in that it can directly analyse the secondary structure of proteins in molecular dynamics trajectories (and thus allows observing changes of secondary structures in time), has a native hydrogen analysis mode, has an alternative to the direct search for neighbors of residues, has the ability to determine the hydrogen bond between residues using a geometric criterion, has the ability to generate data for plotting the number of secondary structures and has no restrictions on the size of the amino acid sequence under consideration. It is shown that the developed algorithm works much faster than the original implementation, but at the same time, it produces identical results. Many additional parameters make the developed algorithm very flexible and customizable.

The data on the secondary structure obtained using the developed DSSP algorithm on the molecular dynamics trajectory of the defensin-like protein of *Pentadiplandra brazzeana* correspond not only to the data obtained using the DSSP v.4 algorithm, but also to the experimental data: all obtained values of the secondary structure are in accordance with the protein folds in the crystal structures.

## Data Availability

Implementation of our algorithm available in Gromacs distribution since version 2023. Also we created standalone version of DSSP algorithm implementation in Gromacs is available at the GitLab repository: https://gitlab.com/bio-pnpi/gmx-dssp. You can also compare results obtained with DSSP v.4 with results obtained with our implementation using the script available at https://gitlab.com/bio-pnpi/gmx-dssp-refdata.

## Acknowledgments

The enhanced DSSP algorithm was developed and implemented within the state assignment of Ministry of Science and Higher Education of the Russian Federation (theme N^*o*^ 121060200127-6 and theme N^*o*^ 2.8629.2017). The structures and molecular dynamics trajectories of scFv antibodies were obtained within the state assignment of Ministry of Science and Higher Education of the Russian Federation (theme N^*o*^ 075-15-2021-1360).

## Supporting Information Available

### Initial structures

Initial structures for test objects were taken from the following sources:

- structure of the defensin-like protein of *Pentadiplandra brazzeana* was taken from PDB ID 1BRZ
- structure of NSP12 from SARS-Cov-2 was taken from PDB ID 7BTF
- structure of T7 polymerase was taken from PDB ID 2PI5

Test objects 3B12, 3H10, B11-1, B11-2 and ST4 are scFv antibody. Seq sequences used to create them were constructed in a following way. Briefly, mice were immunized with recombinant VEGFR-1, CTLA-4 and Stabilin-1 proteins. Obtained B-lymphocytes were fused to myeloma cells and producing monoclonal antibodies hybridomas: 3B12 and 3H10 variants against VEGFR-1, B11-1 and B11-2 against CTLA-4 and St4 against Stabilin-1, were selected. The genes of the variable parts of heavy and light chains (VH and VL respectively) were sequenced and converted into amino acid sequences. Next, VH and VL fragments were connected into a single polypeptide chain via a (G4S)3 flexible linker in a «head-to-tail» manner, and tagged with a GSS-His6 at the C terminus. Thus, the final domain structure of these scFv antibodies is VH-(G4S)3-VL-GSS-His6. Each scFv antibody structure was modelled using AlphaFold.^11^

### Molecular dynamics

Molecular dynamics (MD) modeling was performed with the software package GROMACS 2023,^12^ the amber14sb field^13^ was used for the protein and tip3p was used as a water model. The resulting systems were placed in a periodic water box in such a way that at least 25Å remained to the walls of the box. Then the resulting box was minimized using the steepest descent algorithm. The system obtained as a result of minimization was subjected to charge neutralization by adding 150 mM of NaCl bringing the total charge of the system to zero. The neutralized system was again subjected to the procedure of energy minimization using the steepest descent algorithm. Next, the system was equilibrated using a two-stage approach. At the first stage, all heavy atoms of protein were restrained to their initial positions using an additional energy term (posres), while at the start of equilibration, the temperature (particle velocity distribution) was taken from the Maxwell distribution for a given temperature at 310K. The system was equilibrated for 5 ns at each temperature. The integration step was 2 fs; the V-rescale thermostat and the C-rescale barostat were used. At the second stage, the additional restraining potential was removed, and all components of the system could move freely. During this stage, the Nose-Hoover thermostat and the Parrinello-Rahman barostat were used, and the system was equilibrated for 10 ns. The final state obtained as a result of a two-stage equilibration was used as a start for the production dynamics. Molecular dynamics was carried out for 1000 ns, using the same set of parameters as for the second stage of equilibration.

